# Corticostriatal Responses to Social Reward are Linked to Trait Reward Sensitivity and Subclinical Substance Use in Young Adults

**DOI:** 10.1101/2023.01.17.524305

**Authors:** James B. Wyngaarden, Camille R. Johnston, Daniel Sazhin, Jeff B. Dennison, Ori Zaff, Dominic Fareri, Michael McCloskey, Lauren B. Alloy, David V. Smith, Johanna M. Jarcho

**Author notes:** **Correspondence** David V. Smith or Johanna M. Jarcho. denotes co-first authorship (James B. Wyngaarden, Camille R. Johnston). denotes co-senior authorship (David V. Smith, Johanna M. Jarcho).

## Abstract

Aberrant levels of reward sensitivity have been linked to substance use disorder and are characterized by alterations in reward processing in the ventral striatum (VS). Less is known about how reward sensitivity and subclinical substance use relate to striatal function during social rewards (e.g., positive peer feedback). Testing this relation is critical for predicting risk for development of substance use disorder. In this pre-registered study, participants (N=44) underwent fMRI while completing well-matched tasks that assess neural response to reward in social and monetary domains. Contrary to our hypotheses, aberrant reward sensitivity blunted the relationship between substance use and striatal activation during receipt of rewards, regardless of domain. Moreover, exploratory whole-brain analyses showed unique relations between substance use and social rewards in temporoparietal junction. Psychophysiological interactions demonstrated that aberrant reward sensitivity is associated with increased connectivity between the VS and ventromedial prefrontal cortex during social rewards. Finally, we found that substance use was associated with decreased connectivity between the VS and dorsomedial prefrontal cortex for social rewards, independent of reward sensitivity. These findings demonstrate nuanced relations between reward sensitivity and substance use, even among those without substance use disorder, and suggest altered reward-related engagement of cortico-VS responses as potential predictors of developing disordered behavior.

## Introduction

From April 2020 to April 2021, over 92,000 people died from drug overdose in the United States—the highest death toll during a 12-month period on record (NCDAS, 2022). Today, over 20 million Americans are affected by substance use disorder (Avena et al., 2021), and as such, understanding the risk factors associated with problematic substance use is a primary public health concern. Understanding individual differences in reward processing may provide insight into substance use and shed light on individuals most at risk for developing substance use disorder down the road (e.g., Bart et al., 2021). Trait reward sensitivity is a self-reported predisposition to seek rewarding substances and experiences (e.g., food, sex, etc.; Volkow et al., 2010). Neuroimaging studies have shown that reward sensitivity is associated with activity in the ventral striatum (VS) and medial prefrontal cortex (Beaver et al., 2006; Simon et al., 2010; Sripada et al., 2011; Pujara et al., 2016). One theory of risk for substance use disorder argues that repeated substance use over time blunts reward-related activation in the VS and ventromedial prefrontal cortex (vmPFC; Beck at al., 2009; Goldstein & Volkow, 2011). Thus, individuals with a hyposensitive reward system may be predisposed toward using psychoactive substances to compensate and up-regulate their reward systems (Blum et al., 1996). Other evidence suggests substance use is associated with hypersensitivity to rewards. For example, higher behavioral approach scores, which are closely linked to reward sensitivity, are associated with alcohol use (Franken & Muris, 2006), cocaine addiction (Balconi & Finocchiaro, 2015), and nicotine dependence (Lawn et al., 2015). Moreover, hypersensitive individuals tend to be more impulsive and risk-seeking, factors that also may contribute to substance use (Nusslock & Alloy, 2017). Thus, individuals with high or low (i.e., aberrant) reward sensitivity are at risk for developing substance use problems, irrespective of their current expression of substance use. These seemingly conflicting observations suggest that disparate mechanisms may lead to a common clinical outcome, and they underscore the importance of examining individual differences in trait reward sensitivity. However, these findings are also limited in their consideration of external forces, like social factors, and how they contribute to risk for developing substance use problems.

While substance use often occurs in a social context (Goncy et al., 2013; Wei et al., 2010; Andrews et al., 2002; Dishion & Owen, 2002) and social stressors, such as peer rejection, commonly precipitate consumption (Nepon et al., 2021; Milivojevic & Sinha, 2018), relationships between substance use and reward sensitivity in the context of *social* rewards are surprisingly understudied. Even fewer studies include brain responses to both social and monetary rewards, which are necessary to make claims about the specificity of effects. Additionally, associations between substance use and VS-based connectivity during receipt of social rewards remain relatively unexplored. Social rewards elicit VS activation (Izuma et al., 2008), and recent meta-analyses show that processing social information is related to engagement of the dorsal and ventral medial prefrontal cortex (dmPFC, vmPFC), posterior cingulate cortex (PCC), right fusiform face area (FFA), and bilateral amygdala (Tso et al., 2018; Feng et al., 2021; Martins et al., 2021). Therefore, such regions may be promising targets for VS connectivity during receipt of social rewards (e.g., Sazhin et al., 2020; Dennison et al., 2023). When assessing these relationships, controlling for trait reward sensitivity is important, as doing so helps to disentangle the specific effects of substance use behaviors from individual differences in reward responsiveness, a risk factor for substance use disorder.

Adolescence and emerging adulthood are periods of heightened sensitivity to substance use, characterized by increased risk for and expression of substance use disorder (Crews et al., 2007; Wood et al., 2018), particularly given that adolescents with aberrant reward sensitivity scores are more likely to develop substance use disorder (Alloy et al., 2009; Ivanov et al., 2008; Genovese & Wallace, 2007). Importantly, this developmental stage coincides with profoundly heightened neural sensitivity to social rewards, like peer approval (Foulkes & Blakemore, 2016; Nelson et al., 2016; Somerville, 2013). Despite these insights, understanding the brain-based relationship between substance use and reward sensitivity may be limited by a reliance on narrowly focused reward paradigms (e.g., only evaluating monetary outcomes; e.g., Dennison et al., 2022; Bart et al., 2021) performed by individuals who have already developed substance use disorder. Studying diverse reward domains, including social feedback, in a population who is at risk for, but yet to express substance use disorder is critical to differentiate between causal factors and consequences associated with problematic substance use behaviors. If an individual is already using substances heavily, then it is hard to distinguish their current usage patterns from their future risk of having problems. Similarly, abnormal brain function may precede problematic substance use, or it may emerge due to problematic substance use behavior. To this end, efforts focused on prevention of substance use problems must target healthy individuals who are at risk for developing problems.

We leveraged well-matched social and monetary reward tasks (Distefano et al., 2018; Nelson & Jarcho, 2021; Quarmley et al., 2019) to investigate associations between reward sensitivity, subclinical substance use, and brain activation and connectivity. First, we aimed to quantify the relationship between reward sensitivity, substance use, and VS activation during social compared to monetary rewards. Our hypotheses were informed by prior work showing that that substance use frequently takes place in social settings and is often triggered by social stressors such as peer rejection, suggesting that individuals with greater substance use may be especially attuned to rewards in the social domain. Thus, we hypothesized that individuals with higher levels of substance use would show exaggerated VS responses to social vs. monetary rewards, independent of reward sensitivity. Second, we aimed to examine the relationship between reward sensitivity, substance use, and VS connectivity during social vs. monetary rewards. We hypothesized that individuals with higher levels of substance use would show enhanced connectivity between the VS and regions modulated by social information, independent of reward sensitivity. However, we also conducted a series of exploratory analyses including the interaction between substance use and reward sensitivity to assess the role of reward sensitivity as a moderator of risk for future substance use problems.

## Materials and Methods

### Participants

Although the pre-registration (https://aspredicted.org/blind.php?x=JNH_EGK) describes our goal to collect data from 100 participants (18-22), we ultimately studied 59 participants due to constraints imposed by the COVID-19 pandemic. Using pre-registered criteria, fifteen of the 59 participants who completed the study were excluded from analyses due either to failure to respond during behavioral tasks (>20% missing responses; N=4), incomplete data (N=5; failure to complete survey data or missing behavioral data due to technical issues), or poor image quality (N=6). Image quality was defined using the fd_mean and tSNR values from MRIQC (Esteban et al., 2017). Participants were excluded for fd_mean values greater than 1.5 times the interquartile range, or for tSNR values below the lower bound of 1.5 times the interquartile range, per the distribution from neuroimaging data of otherwise eligible participants. This resulted in a final sample of 44 participants (mean age: 20.45 years, SD: 1.89 years; 22.7% male; 57% white, 34% Asian, 9% other—2 Black/African American, 1 Black and white, 1 Indian).

Participants were recruited via the Temple University Psychology and Neuroscience Department participant pool, and from the surrounding community via flyers and online advertisements. Participants were paid $25 per hour for fMRI and $15 per hour for behavioral tasks, and received bonuses based on their decisions on other neuroeconomic tasks (not reported here), resulting in a total payment of $140 to $155. In addition, participants recruited from the university pool also received research credit for their participation.

### Procedure

All methods were approved by the Temple University IRB. Prospective participants were identified based on their responses to an online screener, which assessed reward sensitivity using the Behavioral Activation Subscale (BAS; Carver & White, 1994) and the Sensitivity to Reward subscale (SR; Torrubia & Tobeña, 1984). A sum was calculated for each subscale. Sums were assigned to a quintile that reflected low to high levels of reward sensitivity across the distribution of possible scores. Using methods consistent with our prior work (e.g., Alloy et al., 2009) designed to increase confidence that participants were truthful, attentive, and not responding at random, only participants with scores within +/-1 quintile on both subscales were eligible for the study (no exclusions were made based on this criteria). Substance use information also was collected on the online screener via the Alcohol Use Disorders Identification Test (AUDIT; Babor et al., 1992) and the Drug Use Identification Test (DUDIT; Berman et al., 2002). At the in-person visit, we confirmed that eligible participants were free of major psychiatric or neurologic illness and MRI contraindications. Breathalyzer tests, urine drug screens, and (for females) urine pregnancy tests were also collected to confirm the absence of alcohol, illicit drugs, and pregnancy prior to fMRI.

Participants were told that they would complete a social evaluation task with peers who had already completed the study. Prior to their fMRI visit, our team took a photograph of the participant that was purportedly uploaded to a study database. Participants believed that once this photograph was uploaded, peers would receive a text message on their cell phone asking them to view the photo and indicate whether they thought they would ‘like’ or ‘dislike’ the participant. Participants were told that at their fMRI visit several days later, they would be asked to guess which peers ‘liked’ and ‘disliked’ them. Participants were also told that they would be completing a monetary guessing task. Data is available on OpenNeuro (https://openneuro.org/datasets/ds004920/versions/1.1.0; for more detail, see Smith et al., 2024).

#### fMRI-Based Tasks

The monetary and social reward tasks (Distefano et al., 2018; Nelson & Jarcho, 2021; Quarmley et al., 2019) were administered using PsychoPy (Peirce et al., 2019). As depicted in Figure 1, there were two tasks (monetary, social) that were presented in separate runs (order counterbalanced across participants). This design eliminates task-switching costs (Hayden et al., 2010) while still preserving our ability to directly contrast responses associated with social and nonsocial reward. In the monetary task, participants were shown two doors.

**Figure 1.**
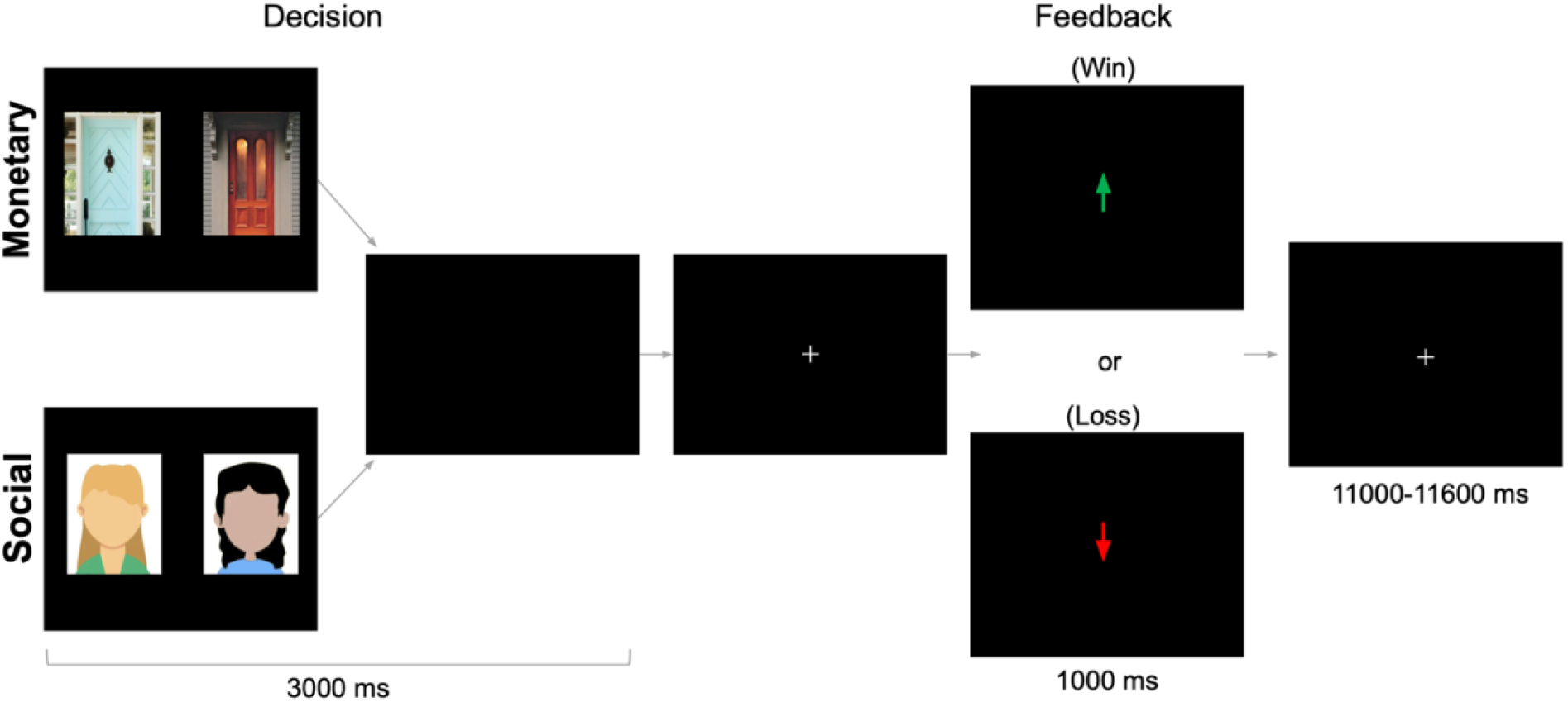
fMRI-based monetary and social reward tasks. On each trial, participants choose either between two doors (monetary task) or the faces of two peers (social task) in search of a reward. After a brief interval, they receive feedback: an upward arrow indicating a win (monetary task=$0.50 gain; social task=positive peer feedback) or a downward arrow indicating a loss (monetary task=$0.25 loss; social task=negative peer feedback). Tasks were administered separately but were matched on these dimensions. Results focus on the outcome period.

Participants were instructed to pick the door that contained a $0.50 prize. On monetary reward trials, feedback indicated that the participant won $0.50. On monetary loss trials, feedback indicated that the participant lost $0.25. In the social task, participants were shown the faces of two peers. Participants were informed that each peer had indicated whether they liked or disliked the participant based on a photo of them, and they were instructed to pick the peer that liked them. On social reward trials, feedback indicated that the peer liked them. On social loss trials, feedback indicated that the peer disliked them (See supplementary materials for more details). Each run included 60 trials: 50% resulted in reward and 50% resulted in loss feedback. Although both tasks involve feedback about correct/incorrect guesses, the result of that feedback occurs in social or monetary domains. Each task collected 292 functional volumes. Trials were separated by a variable duration intertrial interval (1,100–11,600 ms; M = 3,500 ms).

To assess whether participants were deceived by the peer feedback in the social task, we used a series of post-experimental survey items including both direct (e.g., “when you were completing the social task, were others rating you?”) and indirect inquiries (e.g., “When you saw that someone liked you based on your photo, how happy/excited did you feel?”). Based on these responses and other qualitative aspects of the visit, the experimenter rated the extent to which they believed the participant was deceived. Using these criteria, we determined that a subset of participants (n=8) were not deceived during the social reward task. However, these subjects did not differ from deceived participants in their response times for social relative to monetary reward (F=0.140, p=0.708), indicating no difference in levels of engagement across participants. As such, these participants were included in analyses to retain statistical power, consistent with prior studies using this task (e.g., Quarmley et al., 2019; Jarcho et al., 2022).

### Individual Difference Measures

#### Trait Reward Sensitivity

Reward sensitivity was defined by a composite score (*α*=0.718) consisting of the sum of the z-scores for the Behavioral Activation Scale (BAS; Carver & White, 1994) and the Sensitivity to Reward (SR) subscale of the Sensitivity to Punishment/Sensitivity to Reward Questionnaire (Torrubia et al., 2001). The BAS is a 20-item self-report questionnaire that aims to measure sensitivity to appetitive motives (e.g., “I go out of my way to get things I want”). The SR is a 24-item self-report measure aimed at more context-specific items. The total BAS and the SR subscale are reliable and valid measures of reward sensitivity (Alloy et al., 2006; Alloy et al., 2012).

We tested both linear and nonlinear associations between brain responses and reward sensitivity. However, assessing nonlinear associations with reward sensitivity (i.e., aberrant reward sensitivity) by taking the 2^nd^-order polynomial expansion overweights the tails of the distribution (Buchel et al., 1998). To avoid this, we first normalized the values by converting them to deciles, where each decile had approximately the same number of participants (cf. Winsorizing). Although this strategy was not described in our pre-registration, we believe it is a necessary deviation because it ensures that our analyses can account for both linear and nonlinear associations (e.g., U-shaped and inverted U-shaped patterns between brain responses and reward sensitivity) while ensuring that results are not driven by extreme values in the tails of the distribution.

#### Substance Use

To distinguish risk factors from consequences of problematic substance use, it is crucial to study individuals who have not yet exhibited problematic substance use. Current substance use was defined by a composite score consisting of the sum of the z-scores for the Alcohol Use Disorders Identification Test (AUDIT; Babor et al., 1992) and the Drug Use Identification Test (DUDIT; Berman et al., 2002). The AUDIT is a 10-item self-report measure that assesses frequency (e.g., “How often do you have a drink containing alcohol?”) and disruptiveness (e.g., “How often during the last year have you failed to do what was normally expected of you because of drinking?”) of alcohol use. Scores greater than 15 are categorized as high risk for alcohol dependence (Babor et al., 2001). No participant scored in the high-risk category for AUDIT (M=3.62, SD=3.53). The DUDIT is an 11-item self-report measure that assesses frequency and disruptiveness of non-alcoholic drug use, containing references to a wide array of substances, including marijuana, cocaine, and others. Scores greater than 24 are categorized as high-risk for dependence (Berman et al., 2005). Only one participant scored in the high-risk category for DUDIT (score=26; M=1.65, SD=4.30). These scores on the AUDIT and DUDIT suggest that our sample exhibits relatively less substance use than what is typical for this age group (Blows & Isaacs, 2022; Shorey et al., 2013; Verhoog et al., 2020). Due to the low scores across measures, we characterize substance use in our sample as “subclinical.” We summed z-scores for AUDIT and DUDIT because we did not have hypotheses differentiating between alcohol and drug use.

### Neuroimaging Data Acquisition and Preprocessing

Functional images were acquired using a 3.0 Tesla Siemens PRISMA MRI scanner and a 20-channel head coil. Neuroimaging data were converted to the Brain Imaging Data Structure (BIDS) using HeuDiConv version 0.9.0 (Halchenko et al., 2019). Results included in this manuscript were preprocessed with fMRIPrep 20.2.3 (Esteban et al., 2018a, 2018b), which is based on Nipype 1.4.2 (Gorgolewski et al., 2011, 2018). MRI acquisition and preprocessing parameters are described in the supplementary materials.

### Neuroimaging Analyses

#### Individual Level Analyses

Neuroimaging analyses used FSL version 6.0.4 (Smith et al., 2004; Jenkinson et al., 2012). We conducted two types of analyses (activation and connectivity) to investigate how social, compared to monetary, rewards and substance use were associated with BOLD responses, independent of linear or quadratic expressions of reward sensitivity. Exploratory analyses also examined reward sensitivity as a covariate of interest. Both used individual level general linear models with local autocorrelation (Woolrich et al., 2001).

The first model focused on the brain activation evoked during the feedback phase of each reward task (i.e., monetary and social) and used four task-based regressors. Two regressors of interest included reward and loss feedback (duration = 1,000 ms). Two regressors of no interest included the decision phase (duration = 3,000 ms) and trials with missed responses within the decision phase (i.e., failures to respond; duration = 3,000 ms).

The second model focused on task-dependent connectivity with VS as it related to the varying types of feedback in the task (i.e., reward vs. loss). To estimate the changes in connectivity between feedback types, we used psychophysiological interaction (PPI) analysis (Friston et al., 1997; O’Reilly et al., 2012), which can reveal consistent and specific task-dependent changes in connectivity (Smith et al., 2016; Smith & Delgado, 2017). We focused on feedback-dependent changes in connectivity with the bilateral VS (defined by the Oxford-GSK-Imanova atlas; Tziortzi et al., 2011). This model used a total of nine regressors. The first four were identical to those described in the activation model (i.e., reward, loss, decision, and missed trials). A fifth regressor consisted of the average timecourse of activation from the VS seed region (i.e., the physiological regressor). Four additional regressors corresponded to the interaction between the physiological regressor and each of the four original regressors.

Both activation and connectivity models included additional regressors of no interest that controlled for six motion parameters (rotations and translations), the first six aCompCor components explaining the most variance, non-steady state volumes, and framewise displacement (FD) across time. Finally, high-pass filtering (128s cut-off) was achieved using a set of discrete cosine basis functions.

### Hypothesis 1

#### ROI Group Level Analysis

Hypothesis 1 seeks to test whether individuals reporting higher levels of substance use show exaggerated VS responses to social rewards relative to monetary rewards, independent of expressions of reward sensitivity (i.e., domain x feedback x substance use interaction). To account for potential relations with reward sensitivity, both linear (i.e., greater values correspond with greater reward sensitivity) and quadratic (i.e., greater values correspond with more aberrant reward sensitivity) measures of reward sensitivity were included as regressors. For each participant, activation in the VS seed described above was extracted using AFNI’s 3dROIstats. We used FSL’s PALM (Winkler et al., 2014; Alberton et al., 2020) to conduct a linear regression for the difference in extracted striatal BOLD for social compared to monetary reward processing (see Table 1 for notation) that was regressed onto a model of substance use including additional covariates for reward sensitivity (first- and second-order measures), the substance use x reward sensitivity and substance use x reward sensitivity^2^ interactions, and nuisance regressors (tSNR and motion; seven covariates total). Lower order effects of interest were also probed in the absence of higher order interactions. Given that social and monetary tasks were administered separately, it is critical that we account for differences in the quality of confounds like temporal signal-to-noise ratio (tSNR) and framewise displacement. To account for these differences, we subtracted the value from the monetary task from the social task. Given signal was extracted signal from a single, a priori ROI, correction for multiple comparisons was not required to test this hypothesis (for further discussion, see Gentili et al., 2020 and Rubin, 2021).

**Table 1.** Terms for model contrasts.

| Term | Effect | Contrast |
| --- | --- | --- |
| Social reward processing | Domain x Feedback | Social (Reward > Loss) > Monetary (Reward > Loss) |
| Reward processing | Feedback | (Social Reward + Monetary Reward) > (Social Loss + Monetary Loss) |
| Social processing | Domain | (Social Reward + Social Loss) > (Monetary Reward + Monetary Loss) |
| Decision phase | Social Stimuli | (Social Decision > Baseline) > (Monetary Decision > Baseline) |

### Hypothesis 1 Exploratory Analyses

#### ROI Group Level Analysis

We hypothesized that we would observe a domain x feedback x substance use interaction, such that individuals reporting higher levels of substance use would show exaggerated striatal responses to social rewards relative to monetary rewards, independent of reward sensitivity. However, lower order interactions that included task effects (i.e., domain or feedback) were also explored to test the extent to which substance use may relate to brain activity as a function of other task features. Exploratory ROI-based analyses also examined relations with reward sensitivity as a covariate of interest, in order to assess whether reward sensitivity may serve as an indicator of risk for future substance use problems. These analyses use data extracted from the ventral striatum and thus do not correct for multiple comparisons (e.g., Gentili et al., 2020; Rubin, 2021)

#### Whole Brain Group Level Analyses

We also conducted exploratory whole brain analyses to investigate regions outside of the VS that may be implicated in substance use, reward, and social processes. Group-level analyses were conducted using Randomise (Winkler et al., 2014) and focused on social reward processing using the same linear regression model described above. In addition to substance use, whole brain exploratory analyses also investigated relations with reward sensitivity as a covariate of interest. We note that Z statistic images were thresholded parametrically (Gaussian Random Field Theory) using clusters determined by Z>3.1 and a (corrected) cluster significance threshold of P=.05 (Flandin & Friston, 2019; Nichols & Hayasaka, 2003). These whole brain analyses are corrected for multiple comparisons.

### Hypothesis 2

#### PPI Target Regions

Hypothesis 2 seeks to investigate whether substance use, independent of reward sensitivity, is associated with elevated connectivity between the VS and regions implicated in social information processing (e.g., ventromedial prefrontal cortex) during responses to social, relative to monetary, reward. To this end, we sought to identify brain regions modulated by social information (i.e., stimuli in the social task) in the current dataset. To isolate those regions, we contrasted brain function during the decision phase between the social task, where participants viewed pictures of peers, and the monetary task, where participants viewed pictures of doors. Methodologically, the use of task-based contrasts to define hypothetical targets is consistent with prior work (e.g., Smith et al., 2014; Genevsky et al., 2018), and ensures that the resulting regions are relevant to the task and participants in the current study. Potential targets were constrained by prior research (e.g., Tso et al., 2018; Feng et al., 2021; Martins et al., 2021) and focus only on those included in a priori conceptual models of social processing. Importantly, we note there are significant clusters that emerged from the task-based contrasts that were omitted as potential targets because they are not typically associated with social processing (see https://neurovault.org/images/790566/ for the thresholded zstat image).

Significant clusters of activation were found in several regions. For each potential target region, a 5mm sphere was drawn around the peak voxel in the cluster (see Figure 2 and https://identifiers.org/neurovault.collection:13364). Significant regions included the ventromedial prefrontal cortex (vmPFC; x=33, y=61, z=20), dorsomedial prefrontal cortex (dmPFC; x=34, y=62, z=34), right fusiform face area (rFFA; x=47, y=26, z=20), bilateral amygdala (right, x=39, y=42, z=22; left, x=25, y=42, z=22), and posterior cingulate cortex (PCC; x=33, y=25, z=36).

**Figure 2.**
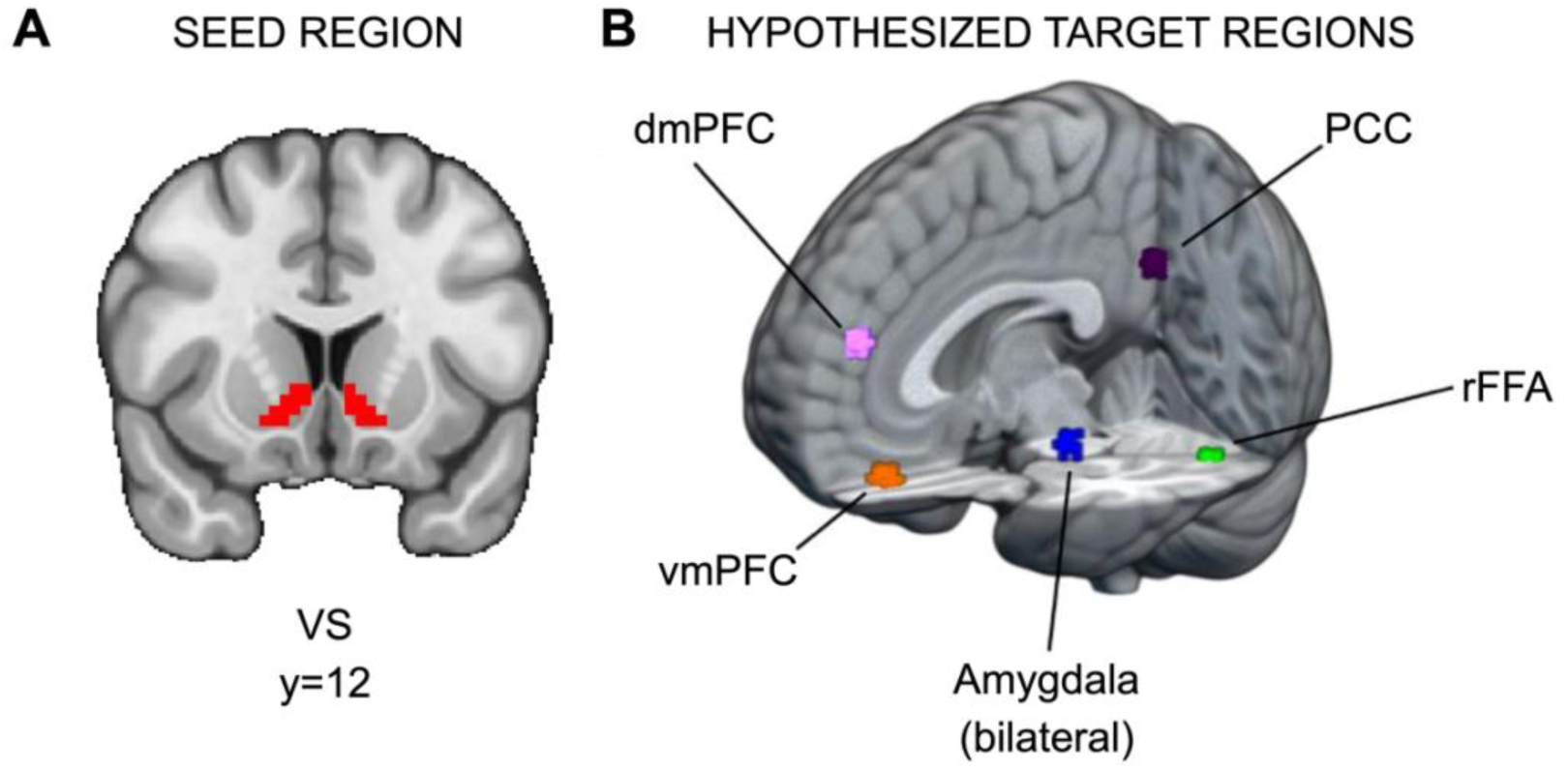
ROIs for PPI analyses. (A) Ventral striatum seed region and (B) hypothesized target regions for PPI analyses (https://identifiers.org/neurovault.collection:13364). Hypothetical target regions related to social information processing in the current dataset were identified via significant clusters in the decision phase. For each target region, a 5mm sphere was drawn around the significant cluster’s peak voxel. The amygdala targets were constrained to voxels within the amygdala as defined by Harvard-Oxford Atlas.

There are two caveats for amygdala: first, for the right amygdala, the cluster extended across multiple regions, so a local maximum within the Harvard-Oxford-defined anatomical region for right amygdala was used instead; second, because amygdala is a smaller anatomical region relative to the other potential targets, after drawing 5mm spheres, both amygdala ROIs were constrained to the Harvard-Oxford anatomical atlas.

#### PPI ROI Analysis

For each participant, connectivity estimates for the target regions described above (vmPFC, dmPFC, rFFA, bilateral amygdala, and PCC) were extracted using AFNI’s 3dROIstats from individual level analyses that modeled the timecourse of VS as a seed. For each target region, we used FSL’s PALM (Winkler et al., 2014; Alberton et al., 2020) to conduct linear regression models for the difference in functional connectivity between VS seed and target ROIs for social reward processing. This was regressed onto a model of substance use including additional covariates for reward sensitivity (first- and second-order measures), the substance use x reward sensitivity and substance use x reward sensitivity^2^ interactions, and nuisance regressors (tSNR and motion). Specifically, we hypothesized that we would observe a domain x feedback x substance use interaction, such that individuals reporting higher levels of substance use would show enhanced corticostriatal connectivity during receipt of social relative to monetary rewards, independent of reward sensitivity. These analyses test multiple comparisons across regions of interest and thus we report family-wise error corrected p values.

### Hypothesis 2 Exploratory Analyses

#### PPI ROI Analysis

Hypothesis 2 examines the domain x feedback x substance use interaction in the hypothesized target regions, with the prediction that individuals reporting higher levels of substance use would show enhanced corticostriatal connectivity during receipt of social rewards, independent of reward sensitivity. However, lower order interactions that included task effects (i.e., domain or feedback) were also explored to test the extent to which substance use may relate to connectivity as a function of other task features. In addition to substance use, exploratory analyses also investigated relations with reward sensitivity. These analyses test multiple comparisons across regions of interest and thus we report family-wise error corrected p values.

## Results

### Hypothesis 1

#### No Association between Substance Use and Ventral Striatum Activation During Social vs. Monetary Rewards

Our first goal was to examine whether higher levels of substance use were associated with exaggerated VS response to social relative to monetary rewards, independent of self-reported reward sensitivity. Inconsistent with our first pre-registered hypothesis, we did not observe a significant association between substance use and striatal activation for social vs. monetary rewards after controlling for reward sensitivity (*t*_(36)_=-1.1077, *p*=0.865; full model results in Supplementary Table 1).

### Hypothesis 1 Exploratory Results

#### Aberrant Reward Sensitivity is Associated with a Diminished Relation between Substance Use and Striatal Activation During Reward Receipt

Exploratory analyses examining the extent to which substance use and reward sensitivity may relate to brain activity as a function of other task features were also conducted. In the absence of higher order interactions, lower order interactions investigating striatal effects related to domain and feedback were assessed. Across all subjects, the difference in striatal BOLD for feedback [(social reward + monetary reward) > (social loss + monetary loss)] showed a significant effect (*t*_(36)_=13.534, *p*<0.001), indicating greater striatal activation for rewards relative to losses (full model results in Supplementary Table 1). Importantly, there was no difference in striatal activation between domains (e.g., *t*_(36)_=-0.774, *p*=0.779), suggesting that degree of reward processing across paradigms was similar. A linear regression for the difference in striatal BOLD for feedback on the interaction between substance use and quadratic (aberrant) reward sensitivity also revealed a significant effect (*t*_(36)_=2.198, *p*=0.018): among individuals in the moderate range of trait reward sensitivity scores, substance use is weakly associated with rewarding feedback, however, among individuals with aberrant trait reward sensitivity, substance use is negatively associated with rewarding feedback (Fig. 3; A mean split on substance use was plotted to facilitate visualization, but all analyses were performed with a continuous measure). A linear regression for the difference in striatal BOLD for domain [(social reward + social loss) > (monetary reward + monetary loss)] revealed no significant relationships.

**Figure 3.**
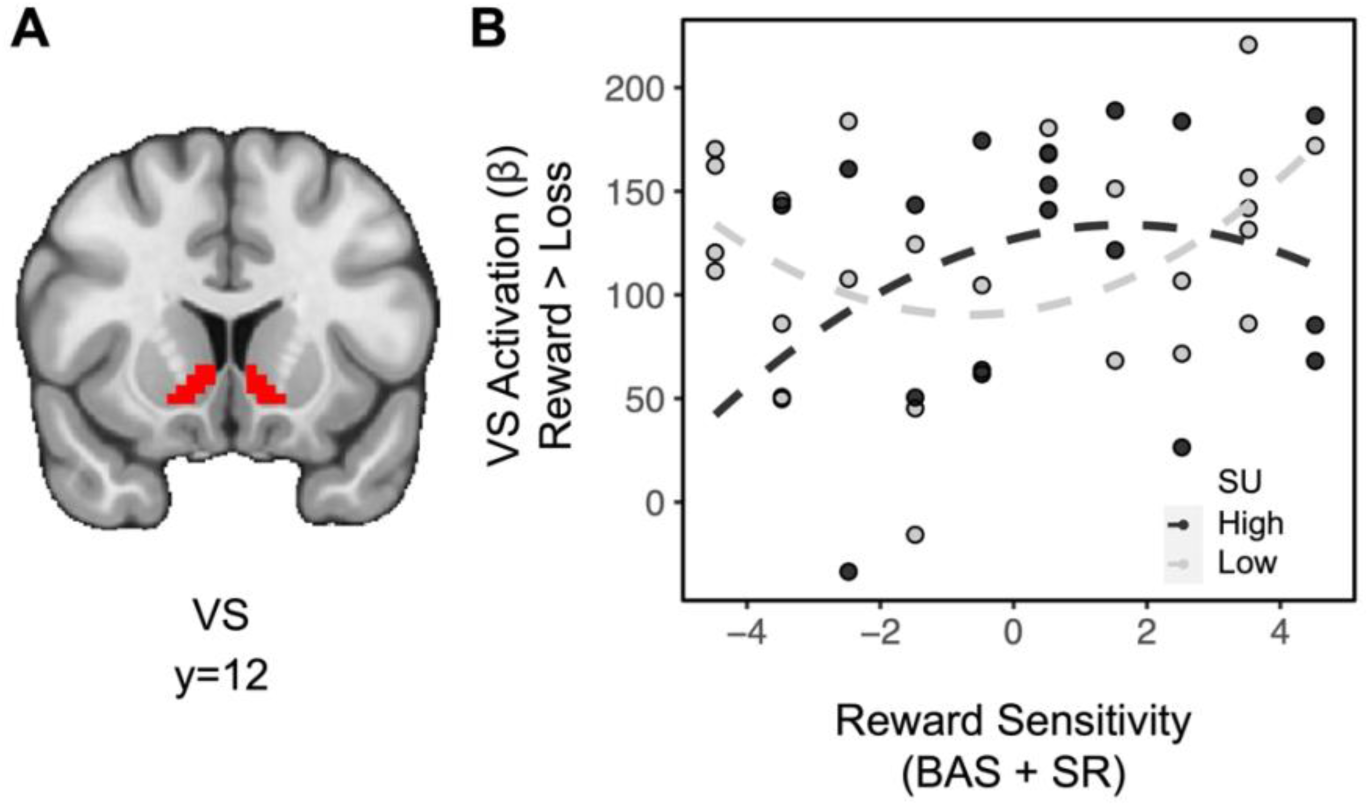
Aberrant reward sensitivity blunts the relationship between substance use and striatal activation during receipt of rewards. (A) Ventral striatum ROI. (B) For individuals with moderate levels of reward sensitivity (near the center of the reward sensitivity measure on the x axis), greater levels of substance use are weakly associated with striatal activation for rewards. However, for individuals with aberrant reward sensitivity (at the extremes of the reward sensitivity measure on the x axis), greater levels of substance use are associated with decreased striatal activation for rewards.

#### Substance Use is Associated with Blunted TPJ Responses to Social vs. Monetary Reward

We also conducted an exploratory whole-brain analysis to investigate the relation between substance use and activation for social rewards beyond the VS. This analysis revealed a significant cluster of activation in the temporoparietal junction (TPJ) for the social rewards in relation to substance use, after controlling for reward sensitivity (see Fig. 4A and https://identifiers.org/neurovault.image:790569). Extracting parameter estimates from the TPJ (x=14, y=19, z=38; ke=56) revealed that as substance use increased, activation in the TPJ decreased for social relative to monetary reward (Fig. 4B).

**Figure 4.**
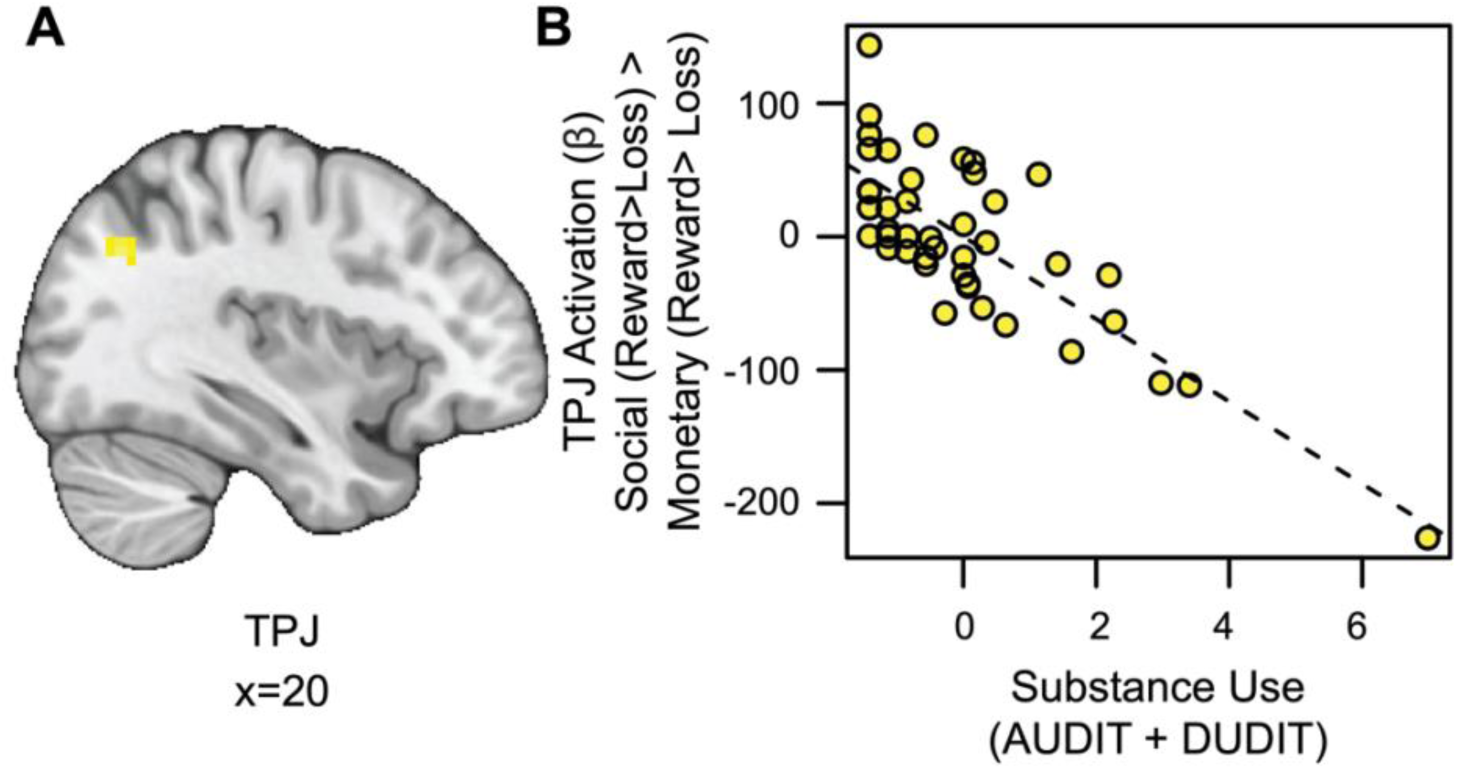
Substance use is associated with blunted TPJ responses to social vs. monetary reward. (A) Cluster-level thresholded activation for social rewards in the temporoparietal junction (TPJ) related to substance use (Thresholded: https://identifiers.org/neurovault.image:790569; Unthresholded: https://identifiers.org/neurovault.image:790568). We note that Z statistic images were thresholded parametrically (Gaussian Random Field Theory) using clusters determined by Z>3.1 and a (corrected) cluster significance threshold of P=.05 (Flandin & Friston, 2019; Nichols & Hayasaka, 2003). (B) While controlling for linear and quadratic measures of reward sensitivity: as substance use increases, social reward activation in the TPJ (component + residual) decreases.

### Hypothesis 2

#### Substance Use is Associated with Reduced Task-Based Connectivity between the VS and dmPFC During Social vs. Monetary Reward

As described in our pre-registration, we hypothesized that substance use, independent of reward sensitivity, would be associated with enhanced striatal connectivity during receipt of social rewards in regions modulated by social information. To test this hypothesis, we conducted an ROI-based psychophysiological interaction (PPI) analysis using the VS as our seed and five social regions as targets. Linear regressions for lower order interactions of domain and feedback also were assessed. No significant effects were observed in the amygdala, rFFA, or PCC for our interactions of interest.

Contrary to our second hypothesis, we found that VS-dmPFC connectivity for social relative to monetary reward was attenuated in individuals with more severe substance use (*t*_(36)_=2.525, *p*=.007, *family-wise-error-corrected p (fwep)*=.0353; full model results in Supplementary Table 2). As substance use increases, connectivity between the VS and dmPFC is reduced (Fig. 5).

**Figure 5.**
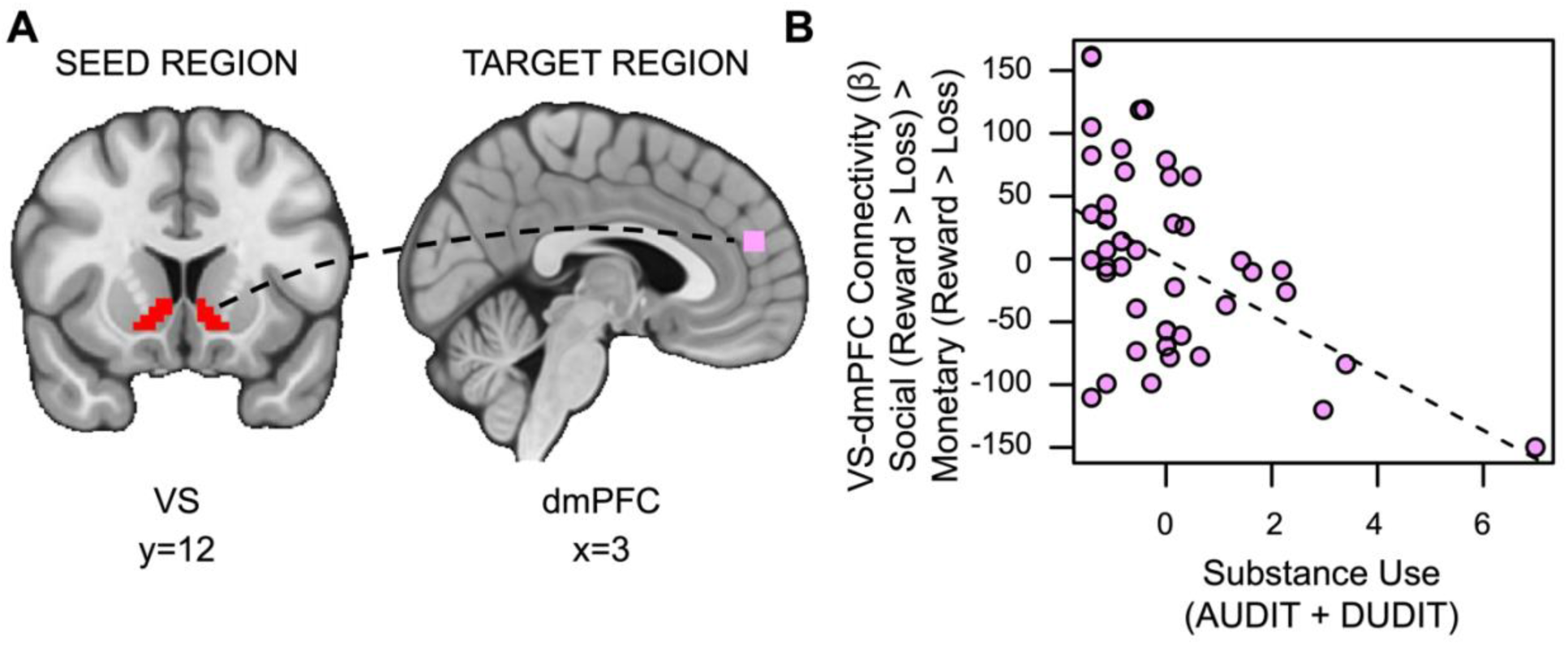
Substance use is associated with decreased VS-dmPFC connectivity for social vs. monetary reward. (A) Ventral striatum seed region and dmPFC target region. (B) As substance use increases, connectivity between the ventral striatum and the dorsomedial prefrontal cortex (dmPFC) during receipt of social rewards is reduced.

### Hypothesis 2 Exploratory Results

Also inconsistent with our hypotheses, social rewards showed enhanced connectivity between the VS and vmPFC in relation to reward sensitivity (*t*_(36)_=2.528, *p*=.007, *fwep*=.0356; full model results in Supplementary Table 3). As reward sensitivity increases, functional connectivity between the VS and vmPFC is enhanced (Fig. 6).

**Figure 6.**
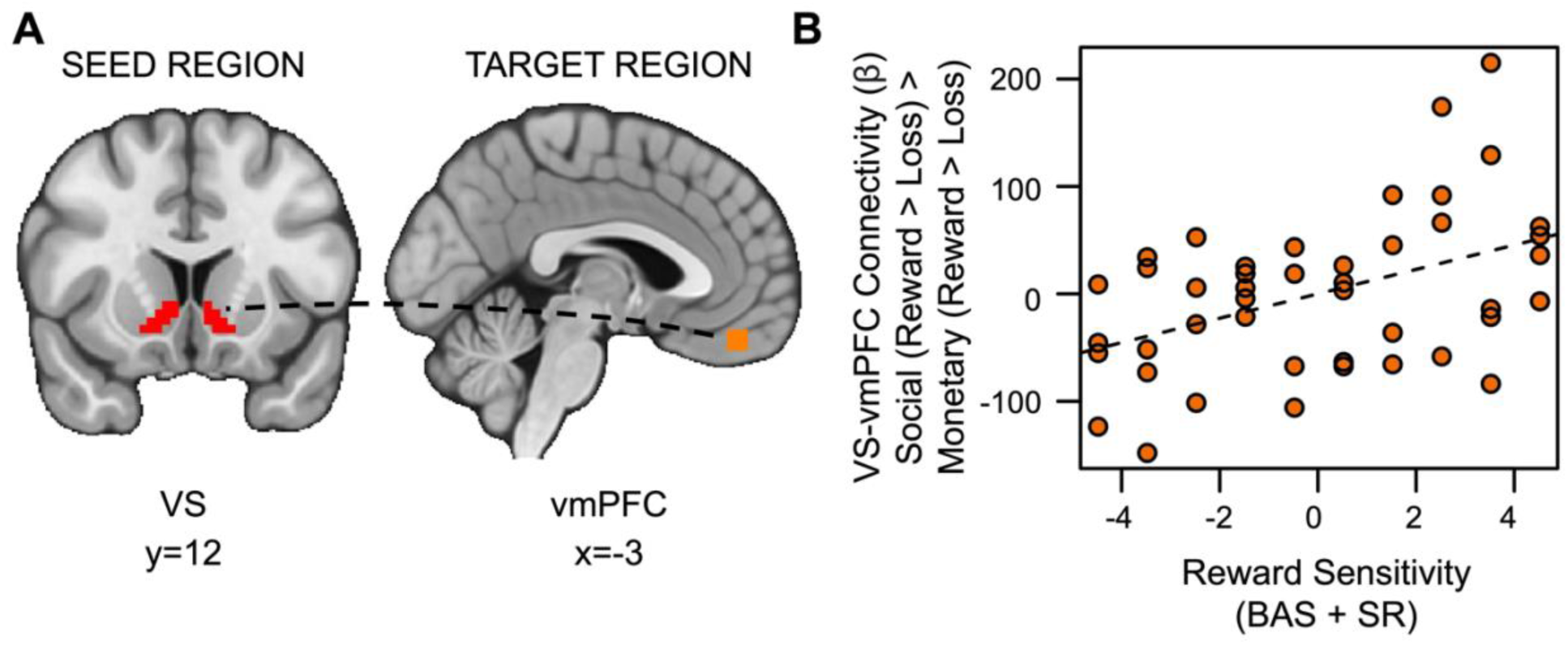
Reward sensitivity is associated with increased VS-vmPFC connectivity for social vs. monetary reward. (A) Ventral striatum seed region and vmPFC target region. (B) As reward sensitivity increases, connectivity between the ventral striatum and the ventromedial prefrontal cortex (vmPFC) during receipt of social rewards is enhanced.

## Discussion

We leveraged well-matched social and monetary reward tasks (Distefano et al., 2018; Nelson & Jarcho, 2021; Quarmley et al., 2019) to investigate associations between reward sensitivity, substance use, and brain activation in young adults with a range of subclinical substance use behavior. Although we hypothesized that substance use would be associated with VS activation for social rewards independent of reward sensitivity, we did not find support for this hypothesis. However, exploratory whole-brain analyses did find this effect in the TPJ, such that substance use was associated with decreased TPJ activation for social vs. monetary rewards while controlling for reward sensitivity. Moreover, exploratory investigation of lower order effects showed that aberrant reward sensitivity blunts the relationship between substance use and VS activation during receipt of domain-general rewards. Contrary to our second hypothesis, our findings show that substance use is associated with decreased VS-dmPFC connectivity for social vs. monetary rewards. Additionally, exploratory analyses showed that reward sensitivity is associated with increased VS-vmPFC connectivity for social vs. monetary rewards. Taken together, these findings suggest that subclinical substance use is associated with blunted response (TPJ) and diminished corticostriatal connectivity (VS-dmPFC) for social relative to monetary rewards, while underscoring the nuanced role of trait reward sensitivity. Moreover, our findings help disentangle results from previous studies that focus on participants with substance use disorder, which make it difficult to determine if altered relations are a cause or consequence of long-term disordered behavior.

Very few studies have investigated the impact of subclinical substance use on processing social rewards (see Beard et al., 2022). However, our finding that substance use is associated with blunted brain activation (TPJ) and connectivity (VS-dmPFC) for social compared to monetary rewards is in line with the handful of recent studies. For example, Zimmermann et al. (2019) found that cannabis users showed less activity in the striatum during interpersonal touch from a female experimenter, while non-users showed more striatal activity. Further, Li et al. (2021) showed that binge drinkers had decreased neural responses in the anterior medial orbitofrontal cortex and precuneus when viewing social interactions of abstract shapes, compared to viewing random movement. Finally, a recent study from our research group (Jarcho et al., 2022), completed after preregistration for the current work, found that greater substance abuse behavior in adolescents was related to decreased right VS response to social rewards (peer ‘like’ feedback). Given that substance use often occurs in a social context (Goncy et al., 2013; Wei et al., 2010; Andrews et al., 2002; Dishion & Owen, 2002) and social stressors commonly precipitate consumption (Nepon et al., 2021; Milivojevic & Sinha, 2018), these domain-specific effects are worth investigating further.

Our findings can be interpreted in various ways. First, substance use was associated with decreased TPJ activation for social vs. monetary rewards, potentially suggesting that increased substance use may be related to deficits in processing social information. Activation in the TPJ is associated with inferring the goals, intentions, and mental states of others (Krall et al., 2015; Van Overwalle 2009) and how they relate to our own (Arioli, et al., 2023). Thus, substance use may blunt one’s ability to process the intentions of others, even – and perhaps especially – during positive social interactions. Second, substance use was related to decreased VS-dmPFC connectivity for social rewards, further reinforcing that increased substance use may be related to altered ability to process social information. Striatal connectivity has been linked to interpreting others’ facial expressions (Kim et al., 2020), and the dmPFC has been implicated in social cognitive functions during receipt of social feedback (e.g., Ferrari et al., 2016), in prediction error during social learning (Joiner et al., 2017; Suzuki et al., 2012), and in emotion regulation/cognitive control (Doré et al., 2017). We speculate that reduced VS-dmPFC connectivity during social reward may reflect impaired facility for assessing and reacting to feedback from others. Alternatively, it may be that those with substance use problems exhibit disrupted regulation of reward processing, particularly in the social domain. More specifically, individuals whose corticostriatal systems are not engaged during rewarding social encounters may subsequently pursue rewarding experiences via alternative means, potentially including substances (Beard et al., 2022).

Additionally, contrary to our hypothesis, trait reward sensitivity was positively associated with VS-vmPFC connectivity for social rewards. Increased subjective utility for a given choice after observing others make a similar choice has been linked to the vmPFC (Chung et al., 2015), and functional connectivity between the VS and the vmPFC has been shown to reflect subjective value for social rewards (Smith et al., 2014; Hill et al., 2017; Hare et al., 2010). The current results suggest that higher trait reward sensitivity is associated with increased functional integration of these regions, perhaps bridging the gap to previous research linking reward sensitivity to social functions like extraversion (Lucas et al., 2000) and anxiety (Booth & Hasking, 2009) which may center on receiving feedback from peers.

There are several future directions to consider when evaluating these results. A benefit of studying those at risk for substance use is that effects observed here are unlikely to be a consequence of long-term use that often occurs in clinical populations. However, it should be mentioned that even among young adults without substance use disorder, substance use scores in our sample tended to be lower than the rates that have been reported elsewhere (e.g., Blows & Isaacs, 2022; Shorey et al., 2013; Verhoog et al., 2020). There may be several reasons for this discrepancy. For example, we excluded individuals who were not willing to forgo the consumption of drugs or alcohol in the 24 hours prior to participation in the study. Additionally, our sample was predominantly female (male n=10; < 23%), and while rates did not differ based on gender in our sample, young adult males generally exhibit higher rates of substance use than females (Brady & Randall, 1999). Previous research has shown the association between responses in the nucleus accumbens/VS during reward anticipation and future problematic substance use and relapse (e.g., Büchel et al., 2017; MacNiven et al., 2018). However, while the limited range of substance use in our sample allows us to examine risk factors for substance use disorder, the subtle effects we observe here may differ from observations of a clinical sample. Moreover, wider variability in substance use may help distinguish between the effects of different types of substances (e.g., alcohol vs. marijuana) that may occur in different settings and have more nuanced relationships with social context and reward sensitivity. To help close the gap between cause and consequence, longitudinal studies could be used to further examine the relationship between subclinical substance use and development of substance use disorder. It should also be mentioned that other psychopathologies could be important to assess in this context. For example, our previous work has shown that processing of social rewards can be associated with depression and anxiety (Quarmley et al., 2019; Nelson & Jarcho, 2021), both of which are linked to substance use disorder. Future investigations should more closely examine these relationships.

We also note that various alternatives to our method of generating ROIs for hypothetical target regions could be defensible. For example, while the method of contrasting the decision phase between domains to define hypothetical target regions is consistent with some of our prior work (e.g., Smith et al., 2014), use of a priori conceptual models, meta-analyses, or atlases are also valid means of defining ROIs. Additionally, while the method of drawing a sphere around a peak voxel to create masks is consistent with prior work (e.g., Smith et al., 2014; Utevsky et al., 2014; Davey et al., 2012; Schneider et al., 2020), there may be valid reasons to use other methods (e.g., anatomical atlases), particularly for the larger ROIs (e.g. dmPFC, vmPFC). Other features of the sample may impact generalizability. For example, the current sample is comprised of college students, who tend to have lower levels of substance use disorder relative to their non-college peers. Differences in social factors related to substance use for college vs. non-college individuals may be relevant to consider as well. The sample is also relatively small, primarily white (57%) and Asian (34%), and predominantly female (77.3%). Gender differences in reward sensitivity have been previously reported (e.g., Swartz et al., 2020), and while substance use disorder is more common in males (Brady & Randall, 1999), this gap is narrowing (Steingrimsson et al., 2012). We note that there are no gender differences for reward sensitivity (*t*=-0.22, *p*=.83) or substance use (*t*=-.10, *p*=.92) in the current sample. As neuroimaging research on sex-based differences in relation to substance use and the brain is mixed (e.g., McHugh et al., 2018), further work is needed in larger samples to test for these important biological differences. Moreover, although a subset of participants were not deceived during the social task, these subjects did not differ significantly in their response times to social reward relative to monetary reward, indicating behavioral engagement. This finding is in line with previous research in which explicitly artificial social feedback has elicited robust neural responses (e.g., Hsu et al., 2015; Sankar et al., 2019; Yttredahl et al., 2018), indicating that deception is not always necessary to elicit social effects. Future research should further explore the interplay between deception and social effects.

Despite these limitations, the present results demonstrate that varying levels of problematic substance use in a subclinical population is associated with variations in activation and task-based connectivity in regions implicated in social processing while experiencing social rewards. Moreover, we show that trait reward sensitivity is an important factor when individuals experience social feedback. Although substance use disorder is a complicated issue, our findings help characterize the important roles in how social factors and aberrant reward sensitivity are related to subclinical substance use. These results may contribute to our understanding of how to identify and reduce instances of substance use disorder in the future.

## Acknowledgements

This work was supported in part by grants from the National Institute of Mental Health (R01-MH123473 and R01-MH126911 to LBA, R01-MH132727 to JMJ), the Eunice Kennedy Shriver National Institute of Child Health and Human Development (R21-HD093912 to JMJ), the National Institute on Aging (RF1-AG067011 to DVS), and the National Institute on Drug Abuse (R03-DA046733 to DVS), and also a fellowship from the Temple Public Policy Lab (to JMJ).

## Data and code availability

Analysis code related to this project can be found on GitHub (https://github.com/DVS-Lab/istart-socdoors). In addition, all data will be made available on OpenNeuro before publication.

