## Supplementary Materials for "Corticostriatal Responses to Social Reward are Linked to Trait Reward Sensitivity and Subclinical Substance Use in Young Adults"

### Supplementary Methods 1. Monetary and Social fMRI-based Tasks

**Monetary Reward Task.** Participants were instructed to choose the door behind which there was a $0.50 prize, and were told that if they chose incorrectly, they would receive a $0.25 loss. Each trial began with the presentation of two doors. Participants used a button box to select either the left or right door on the screen. This "decision phase" totaled 3,000 ms. Specifically, when a button was pressed, a white border appeared around the selected door for 500 ms before stimulus offset. After stimulus offset, a blank screen appeared for the remainder of the 3,000 ms decision phase. A fixation cross then was presented (540 ms) before participants received feedback about the accuracy of their choice (1,000 ms). On reward trials, feedback was a green arrow pointing upwards, meaning the participant correctly selected the winning door. On loss trials, feedback was a red arrow pointing downwards, meaning the participant incorrectly selected the losing door. On trials where the participant failed to respond, feedback was randomly selected. Note that the first 12 participants, collected prior to the pandemic, saw pairs of fractals during this task. After data collection resumed, pictures of doors were used instead to more closely match with the realistic features of peer images. This change did not impact response time following monetary rewards vs. losses (Welch Two Sample t-test; $t = 0.75$, $p = 0.46$) nor striatal response to monetary rewards vs. losses (Welch Two Sample t-test; $t = 0.13$, $p = 0.90$). We therefore collapsed across participants with fractal vs. door stimuli.

**Social Reward Task.** The social task was identical to the monetary task, except images of gender-matched (i.e., two female faces or two male faces) and age-matched peers were presented instead of doors. The social reward task consisted of 120 images compiled from multiple sources (internet databases of non-copyrighted images of college-aged individuals). The pictures of purported peers had positive facial expressions, were cropped so that individuals were pictured from their shoulders up and were edited to have an identical solid gray background. Smiling faces were used because they are common in social reward tasks (Richards et al., 2013; Jarcho et al., 2015; Distefano et al., 2018), and are subject to less misinterpretation than neutral faces (Rapee and Heimberg, 1997; Davis et al., 2016). Images were constrained to a standard size (aspect ratio: 11.2 width, 17.14 height). There were an equal number of trials with male and female peers across the reward and loss conditions (30 pairs each, 60 total). Participants were instructed to choose the peer that liked them based on the photograph of the participant. On reward trials, feedback was a green arrow pointing upwards, meaning the participant correctly selected the person who said they would like the participant. On loss trials, feedback was a red arrow pointing downwards, meaning they incorrectly selected the person who said they would dislike the participant. On trials where the participant failed to respond, feedback was randomly selected.

### Supplementary Methods 2. Neuroimaging Data Acquisition and Preprocessing

**Neuroimaging Data Acquisition.** Bold Oxygenation Level-Dependent (BOLD) sensitive functional images were acquired using a simultaneous multislice (multi-band factor = 2) gradient echo-planar imaging (EPI) sequence (240 mm in FOV, TR = 1,750 ms, TE = 29 ms, voxel size of 3.0 x 3.0 x 3.0 $mm^3$, flip angle = 74°, interleaved slice acquisition, with 52 axial slices). Each run included 292 functional volumes. We also collected single-band reference images with each functional run of multi-band data to improve motion correction and registration. To facilitate anatomical localization and co-registration of functional data, a high-resolution structural scan was acquired (sagittal plane) with a T1-weighted magnetization=prepared rapid acquisition gradient echo (MPRAGE) sequence (224 mm in FOV, TR = 2,400 ms, TE = 2.17 ms, voxel size of 1.0 x 1.0 x 1.0 $mm^3$, flip angle 8°). In addition, we also collected a B0 fieldmap to unwarp and undistort functional images (TR: 645 ms; TE1: 4.92 ms; TE2: 7.38 ms; matrix 74×74; voxel size: 2.97×2.97×2.80 mm; 58 slices, with 15% gap; flip angle: 60°).

##### *Preprocessing of Neuroimaging Data*

The following information is adapted from the fMRIPrep preprocessing details; extraneous details were omitted for clarity.

**Anatomical data preprocessing.** The T1-weighted (T1w) image was corrected for intensity non-uniformity (INU) with `N4BiasFieldCorrection`, distributed with ANTs 2.3.3, and used as T1w-reference throughout the workflow. The T1w-reference was then skull-stripped with a *Nipype* implementation of the `antsBrainExtraction.sh` workflow (from ANTs), using OASIS30ANTs as target template. Brain tissue segmentation of cerebrospinal fluid (CSF), white-matter (WM), and gray-matter (GM) was performed on the brain-extracted T1w using `fast` (FSL 5.0.9). Volume-based spatial normalization to one standard space (MNI152NLin2009cAsym) was performed through nonlinear registration with `antsRegistration` (ANTs 2.3.3), using brain-extracted versions of both T1w reference and the T1w template. The following template was selected for spatial normalization: *ICBM 152 Nonlinear Asymmetrical template version 2009c* (TemplateFlow ID: MNI152NLin2009cAsym)

**Functional data preprocessing.** For each of the BOLD runs per subject, the following preprocessing steps were performed. First, a reference volume and its skull-stripped version were generated by aligning and averaging 1 single-band references (SBRefs). A B0-nonuniformity map (or *fieldmap*) was estimated based on a phase-difference map calculated with a dual-echo GRE (gradient-recall echo) sequence, processed with a custom workflow of *SDCFlows* inspired by the `epidewarp.fsl` script (http://www.nmr.mgh.harvard.edu/~greve/fbirn/b0/epidewarp.fsl) and further improvements in HCP Pipelines. The *fieldmap* was then co-registered to the target EPI (echo-planar imaging) reference run and converted to a displacements field map (amenable to registration tools such as ANTs) with FSL's `fugue` and other *SDCflows* tools. Based on the estimated susceptibility distortion, a corrected EPI (echo-planar imaging) reference was calculated for a more accurate co-registration with the anatomical reference. The BOLD reference was then co-registered to the T1w reference using `flirt` (FSL 5.0.9) with the boundary-based registration cost-function. Co-registration was configured with nine degrees of freedom to account for distortions remaining in the BOLD reference. Head-motion parameters with respect to the BOLD reference (transformation matrices, and six corresponding rotation and translation parameters) are estimated before any spatiotemporal filtering using `mcflirt`.

BOLD runs were slice-time corrected using `3dTshift` from AFNI 20160207. First, a reference volume and its skull-stripped version were generated using a custom methodology of *fMRIPrep*. The BOLD time-series (including slice-timing correction when applied) were resampled onto their original, native space by applying a single, composite transform to correct for head-motion and susceptibility distortions. These resampled BOLD time-series will be referred to as *preprocessed BOLD in original space*, or just *preprocessed BOLD*. The BOLD time-series were resampled into standard space, generating a *preprocessed BOLD run in MNI152NLin2009cAsym space*. First, a reference volume and its skull-stripped version were generated using a custom methodology of *fMRIPrep*. Several confounding time-series were calculated based on the *preprocessed BOLD,* notably including framewise displacement (FD).

Additionally, a set of physiological regressors were extracted to allow for component-based noise correction (*CompCor*). Principal components are estimated after high-pass filtering the *preprocessed BOLD* time-series (using a discrete cosine filter with 128s cut-off) for anatomical component correction (aCompCor). For aCompCor, three probabilistic masks (CSF, WM and combined CSF+WM) are generated in anatomical space. The implementation differs from that of Behzadi et al. in that instead of eroding the masks by 2 pixels on BOLD space, the aCompCor masks are subtracted from a mask of pixels that likely contain a volume fraction of GM. This mask is obtained by thresholding the corresponding partial volume map at 0.05, and it ensures components are not extracted from voxels containing a minimal fraction of GM. Finally, these masks are resampled into BOLD space and binarized by thresholding at 0.99 (as in the original implementation). Components are also calculated separately within the WM and CSF masks. For each CompCor decomposition, the *k* components with the largest singular values are retained, such that the retained components' time series are sufficient to explain 50 percent of variance across the nuisance mask (CSF, WM, combined, or temporal). The remaining components are dropped from consideration. The head-motion estimates calculated in the correction step were also placed within the corresponding confounds file. All resamplings can be performed with *a single interpolation step* by composing all the pertinent transformations (i.e., head-motion transform matrices, susceptibility distortion correction when available, and co-registrations to anatomical and output spaces). Gridded (volumetric) resamplings were performed using `antsApplyTransforms` (ANTs), configured with Lanczos interpolation to minimize the smoothing effects of other kernels.

Many internal operations of *fMRIPrep* use *Nilearn* 0.6.2, mostly within the functional processing workflow. For more details of the pipeline, see the section corresponding to workflows in *fMRIPrep*'s documentation (https://fmriprep.readthedocs.io/en/latest/workflows.html).

Further, we applied spatial smoothing with a 5mm full-width at half-maximum (FWHM) Gaussian kernel using FEAT (FMRI Expert Analysis Tool) Version 6.00, part of FSL (FMRIB's Software Library, www.fmrib.ox.ac.uk/fsl). Non-brain removal using BET (Smith, 2002) and grand mean intensity normalization of the entire 4D dataset by a single multiplicative factor were also applied.

#### Supplementary Table 1. Statistics for regression models of striatal activation

| | Domain x Feedback | | Domain | | Feedback | |
|---|---|---|---|---|---|---|
| Contrast | *tstat* | *p* | *tstat* | *p* | *tstat* | *p* |
| Main Effect + | -0.7739 | 0.7793 | 0.6244 | 0.257 | 13.5344 | 0.0002*** |
| Main Effect - | 0.7739 | 0.2208 | -0.6244 | 0.7431 | -13.5344 | 0.9999 |
| RS + | -0.1752 | 0.5725 | 0.5333 | 0.2985 | 1.5420 | 0.0619 |
| RS - | 0.1752 | 0.4276 | -0.5333 | 0.7016 | -1.5420 | 0.9382 |
| $RS^2$ + | -1.4749 | 0.9226 | 0.0187 | 0.4916 | 0.3478 | 0.3601 |
| $RS^2$ - | 1.4749 | 0.0775 | -0.0187 | 0.5085 | -0.3478 | 0.6400 |
| SU + | -1.1077 | 0.8649 | 0.3464 | 0.3596 | -1.1312 | 0.8693 |
| SU - | 1.1077 | 0.1352 | -0.3464 | 0.6405 | 1.1312 | 0.1308 |
| SU x RS + | -0.2830 | 0.6148 | -0.0643 | 0.5298 | -0.2062 | 0.5794 |
| SU x RS - | 0.2830 | 0.3853 | 0.0643 | 0.4703 | 0.2062 | 0.4207 |
| SU x $RS^2$ + | 0.4043 | 0.3447 | -0.6110 | 0.7281 | -2.1980 | 0.9817 |
| SU x $RS^2$ - | -0.4043 | 0.6554 | 0.6110 | 0.2720 | 2.1980 | 0.0184* |

Note: These findings are based on extracted data from a single, a priori ROI in the ventral striatum.
RS=first-order measure of reward sensitivity; $RS^2$=second-order measure of reward sensitivity (i.e., emphasizes aberrant RS); SU=substance use.
+=positive relationship with striatal activation; -=negative relationship with striatal activation
*(p<.05); **(p<.01); ***(p<.001).

#### Supplementary Table 2. Statistics for regression models of VS connectivity with dmPFC.

| | Domain x Feedback | | | Domain | | | Feedback | | |
|---|---|---|---|---|---|---|---|---|---|
| Contrast | *tstat* | *uncp* | *fwep* | *tstat* | *uncp* | *fwep* | *tstat* | *uncp* | *fwep* |
| Main Effect + | 0.4504 | 0.3200 | 0.7291 | 1.1788 | 0.1249 | 0.4363 | -0.2658 | 0.5976 | 0.922 |
| Main Effect - | -0.4504 | 0.6801 | 0.9672 | -1.1788 | 0.8752 | 0.9999 | 0.2658 | 0.4025 | 0.7813 |
| RS + | 1.7279 | 0.0467* | 0.1757 | -0.0671 | 0.5388 | 0.9590 | -0.9607 | 0.8216 | 0.9924 |
| RS - | -1.7279 | 0.9534 | 0.9999 | 0.0671 | 0.4613 | 0.9367 | 0.9607 | 0.1785 | 0.4735 |
| $RS^2$ + | -0.3105 | 0.6242 | 0.9563 | -0.5649 | 0.7199 | 0.9929 | -0.1298 | 0.5476 | 0.898 |
| $RS^2$ - | 0.3105 | 0.3759 | 0.7923 | 0.5649 | 0.2802 | 0.7646 | 0.1298 | 0.4525 | 0.8316 |
| SU + | -2.5251 | 0.9931 | 1.0000 | 0.3920 | 0.3437 | 0.8307 | 2.1169 | 0.0215* | 0.0868 |
| SU - | 2.5251 | 0.0070** | 0.0353* | -0.3920 | 0.6564 | 0.9836 | -2.1169 | 0.9786 | 1.0000 |
| SU x RS + | 2.0168 | 0.0250* | 0.1017 | 0.9121 | 0.1826 | 0.6021 | 0.4564 | 0.3230 | 0.7135 |
| SU x RS - | -2.0168 | 0.9751 | 1.0000 | -0.9121 | 0.8175 | 0.9993 | -0.4564 | 0.6771 | 0.9586 |
| SU x $RS^2$ + | -0.2995 | 0.6126 | 0.9521 | 0.7860 | 0.2162 | 0.6743 | 0.6934 | 0.2458 | 0.5965 |
| SU x $RS^2$ - | 0.2995 | 0.3875 | 0.8010 | -0.7860 | 0.7839 | 0.9984 | -0.6934 | 0.7543 | 0.9783 |

Note: RS=first-order measure of reward sensitivity; $RS^2$=second-order measure of reward sensitivity (i.e., emphasizes aberrant RS); SU=substance use.
+=positive relationship with VS-dmPFC connectivity; -=negative relationship with VS-dmPFC connectivity
*(p<.05); **(p<.01); ***(p<.001).

#### Supplementary Table 3. Statistics for regression models of VS connectivity with vmPFC.

| | Domain x Feedback | | | Domain | | | Feedback | | |
|---|---|---|---|---|---|---|---|---|---|
| Contrast | *tstat* | *uncp* | *fwep* | *tstat* | *uncp* | *fwep* | *tstat* | *uncp* | *fwep* |
| Main Effect + | 0.0052 | 0.5048 | 0.8795 | 1.8072 | 0.0389* | 0.1716 | -0.1127 | 0.5431 | 0.8886 |
| Main Effect - | -0.0052 | 0.4953 | 0.8891 | -1.8072 | 0.9612 | 1.0000 | 0.1127 | 0.4570 | 0.8338 |
| RS + | 2.5282 | 0.0074** | 0.0356* | 0.6809 | 0.2502 | 0.7327 | 0.4358 | 0.3319 | 0.7108 |
| RS - | -2.5282 | 0.9927 | 1.0000 | -0.6809 | 0.7499 | 0.9968 | -0.4358 | 0.6682 | 0.9587 |
| $RS^2$ + | 0.8768 | 0.1920 | 0.5382 | -1.7859 | 0.9597 | 1.0000 | 0.9296 | 0.1818 | 0.4796 |
| $RS^2$ - | -0.8768 | 0.8081 | 0.9943 | 1.7859 | 0.0404* | 0.1827 | -0.9296 | 0.8183 | 0.9895 |
| SU + | -0.5602 | 0.7031 | 0.9790 | 2.2154 | 0.0153* | 0.0778 | -0.0513 | 0.5201 | 0.8856 |
| SU - | 0.5602 | 0.2970 | 0.7057 | -2.2154 | 0.9848 | 1.0000 | 0.0513 | 0.4800 | 0.8556 |
| SU x RS + | 2.1603 | 0.0190* | 0.0766 | 1.4630 | 0.0758 | 0.3075 | -2.2799 | 0.9847 | 1.0000 |
| SU x RS - | -2.1603 | 0.9811 | 1.0000 | -1.4630 | 0.9243 | 1.0000 | 2.2799 | 0.0154* | 0.0661 |
| SU x $RS^2$ + | 0.8184 | 0.2105 | 0.5700 | -0.4081 | 0.6579 | 0.9910 | 0.0171 | 0.4936 | 0.8672 |
| SU x $RS^2$ - | -0.8184 | 0.7896 | 0.9929 | 0.4081 | 0.3422 | 0.8391 | -0.0171 | 0.5065 | 0.8705 |

Note: RS=first-order measure of reward sensitivity; $RS^2$=second-order measure of reward sensitivity (i.e., emphasizes aberrant RS); SU=substance use.

+=positive relationship with VS-vmPFC connectivity; -=negative relationship with VS-vmPFC connectivity

*(p<.05); **(p<.01); ***(p<.001).

### Supplementary Table 4. Summary of reported results.

| Hyp | Analysis | Effect | Contrast | *tstat* | *p* | *fwep* |
|---|---|---|---|---|---|---|
| H1 | Domain x Feedback | VS act | SU + | -1.1080 | 0.8700 | - |
| H1: ExA | Feedback | VS act | SU x $RS^2$ - | 2.1980 | <.0500* | - |
| H1: ExB | Domain x Feedback | TPJ act | SU - | - | - | <.0500* |
| H2 | Domain x Feedback | VS-dmPFC ppi | SU - | 2.5250 | <.0100** | 0.0400* |
| H2: ExA | Domain x Feedback | VS-vmPFC ppi | RS + | 2.5280 | <.0100** | 0.0400* |

Note: ExA=First exploratory result; ExB=second exploratory result; act=activation; ppi=functional connectivity; RS=first-order measure of reward sensitivity; $RS^2$=second-order measure of reward sensitivity (i.e., emphasizes aberrant RS); SU=substance use; corr-p=corrected p value.
*(p<.05); **(p<.01).
